# No evidence for accelerated brain aging in patients with chronic non-cancer pain

**DOI:** 10.1101/627935

**Authors:** Peter Sörös, Carsten Bantel

## Abstract

Chronic pain is often associated with changes in brain structure and function, and also cognitive deficits. It has been noted that these chronic pain-related alterations may resemble changes found in healthy aging, and thus may represent accelerated or pre-mature aging of the brain. Here we test the hypothesis that patients with chronic non-cancer pain demonstrate accelerated brain aging compared to healthy control subjects. The predicted brain age of 59 patients with chronic pain (mean chronological age ± standard deviation: 53.0 ± 9.0 years; 43 women) and 60 pain-free healthy controls (52.6 ± 9.0 years; 44 women) was determined using the software *brainageR*. This software segments the individual T1-weighted structural MR images into gray and white matter and compares gray and white matter images to a large (n = 2001) training set of structural images, using machine learning. Finally, brain age delta, which is the predicted brain age minus chronological age, was calculated and compared across groups. This study provided no evidence for the hypothesis that chronic pain is associated with accelerated brain aging (Welch’s t-test, p = 0.74, Cohen’s d = 0.061). A Bayesian independent samples t-test indicated moderate evidence in favor of the null hypothesis (BF01 = 4.875, i.e. group means were equal). Our results provide indirect support for recent models of pain related-changes of brain structure, brain function, and cognitive functions. These models postulate network-specific maladaptive plasticity, rather than wide-spread or global neural degeneration, leading to synaptic, dendritic, and neuronal remodeling.

## Introduction

Chronic pain is conceptualized as pain lasting beyond regular tissue healing time without the warning and protection function of physiological nociception (1). Chronic pain is a major health care problem affecting approximately 20% of the adult population (2) and is associated with changes in cognitive and emotional processing (3), such as impairments of executive functions (4), difficulties with attention (5), and number processing (6, 7), as well as an increased risk of depression (2).

While the neurophysiology of nociception is well known, the pathophysiology of chronic pain is far from evident. Acute nociceptive input has been shown to quickly reshape neural circuits on the level of the peripheral and central nervous system, such as the C fiber (8), the spinal dorsal horn (9), and the sensorimotor cortex (10, 11). Molecular, cellular, and systems-level research suggests that maladaptive plasticity in the peripheral and central nociceptive networks contribute to the development of chronic pain (12).

In several types of chronic pain, structural alterations of the brain have been demonstrated, starting with the observation that prefrontal and thalamic gray matter was decreased in chronic back pain patients (13). A decrease of gray matter has also been found in patients with chronic tension type headache (14), migraine (15, 16), episodic cluster headache (17), irritable bowel syndrome (18, 19), osteoarthritis (20), and chronic complex regional pain syndrome (21).

By contrast, other studies found increased gray matter in patients with chronic back pain (in the basal ganglia and thalamus) (22), rheumatoid arthritis (basal ganglia) (23), and chronic vulvar pain (hippocampus, parahippocampal gyrus, and basal ganglia) (24).

In coordinate-based quantitative meta-analyses across various types of chronic pain, several cortical and subcortical areas of decreased and increased gray matter were identified (25, 26). In a smaller number of studies, changes in cerebral white matter have also been investigated (27). In patients with chronic musculoskeletal pain, lower fractional anisotropy, indicating the integrity of white fiber tracts, has been found in the corpus callosum and the cingulum (28).

The functional significance of gray and white matter changes in patients with chronic pain is still unresolved (25). Whether these changes are cause or consequence (29) of chronic pain and whether they represent predisposing factors or compensatory neuroplasticity (30) needs to be determined. Moreover, the neuropathological substrates of these changes are not firmly established. Animal research suggests that rapid changes in synaptic spines and glia cells, as well as neuronal loss and, conversely, neurogenesis may contribute to macroscopic changes in chronic pain (12).

A decrease of total and regional gray matter is one of the hallmarks of brain aging, even in the absence of brain disease (31, 32). Previously, two publications suggested that patients with chronic pain due to fibromyalgia and temporomandibular disorder might experience accelerated or premature brain aging (33, 34). Very recently, a study on elderly (> 60 years) adults concluded that the brains of participants with chronic pain appeared older compared to participants without pain, based on an established machine learning approach (35).

Here we test the hypothesis that patients with chronic non-cancer pain demonstrate accelerated brain aging, when compared to healthy participants, using individual T1-weighted structural MR images and machine learning (36), in a sample of 119 participants between 30 and 68 years of age.

## Methods

### Participants

This study analyzes structural T1-weighted MR images obtained from 119 participants of the ChroPain1 (7) and ChroPain2 studies. The combined sample comprises 59 participants with chronic non-cancer pain and 60 participants free of chronic pain. Patients for ChroPain1 (n = 36) were recruited from the Pain Outpatient Clinic, University Clinic of Anesthesiology, Critical Care, Emergency Medicine, and Pain Management, Klinikum Oldenburg, Oldenburg, Germany. Additional patients were found through advertisements in the local daily newspaper. Patients for ChroPain2 (n = 23) were recruited at a specialized rheumatology practice in Oldenburg, Germany (Dr. Markus Voglau^1^) and through the local ankylosing spondylitis support group. For every patient with chronic pain, a sex- and age-matched (±5 years) pain-free control participant was recruited. Control participants for ChroPain1 (n = 37) and ChroPain2 (n = 23) were identified with the help of the local newspaper, the University of Oldenburg’s web page, flyers, and personal communication.

Inclusion criterion for the pain group was chronic pain, defined as persistent pain for ≥ 12 months. Exclusion criteria for the chronic pain and the healthy control groups were: neurological disorders (such as dementia, Parkinson’s disease, stroke, epilepsy, multiple sclerosis, traumatic brain injury, migraine), psychiatric disorders (such as schizophrenia or major depression), substance abuse, impaired kidney or liver function, and cancer. Since some of the cognitive tests applied in the ChroPain studies depend on cultural background, only participants born and raised in Germany were included. All participants gave written informed consent for participation in one of the studies. A compensation of 10 € per hour was provided. Both studies were approved by the Medical Research Ethics Board, University of Oldenburg, Germany (# 25/2015; 2017-059). ChroPain2 has been pre-registered with the German Clinical Trials Register (DRKS00012791)^2^.

### Magnetic resonance imaging

All MR images were acquired on a research-only Siemens MAGNETOM Prisma (Siemens, Erlangen, Germany) whole-body scanner at 3 Tesla with a 64-channel head/neck coil. The scanner is located at the Neuroimaging Unit, School of Medicine and Health Sciences, University of Oldenburg, Germany. A 3-dimensional T1-weighted MPRAGE sequence was used (37). Imaging parameters for ChroPain1 were: TR: 2000 ms, TE: 2.4 ms, voxel dimensions: 0.7 × 0.7 mm^2^, slice thickness: 0.9 mm, 208 axial slices. Imaging parameters for ChroPain2 were: TR: 2000 ms, TE: 2 ms, voxel dimensions: 0.75 × 0.75 mm^2^, slice thickness: 0.75 mm, 320 axial slices. Both sequences utilized in-plane acceleration (GRAPPA) with a PAD factor of 2 (38). Siemens’ pre-scan normalization filter was used in both sequences for on-line compensation of regional signal inhomogeneities (39).

### Brain age prediction

MR images were first converted from Siemens DICOM format to uncompressed NIfTI (.nii) format using dcm2niix^3^. Structural MR images were then analyzed with the software *brainageR*^4^ (40, 41). For additional methodological details, see the Supplementary Information in Cole et al. 2017 (40) and 2018 (41).

*brainageR* employs SPM12 (University College London, London, UK^5^ for image preprocessing. Using SPM12’s *Segment* function (42), images were bias field corrected and segmented into gray matter, white matter, and cerebrospinal fluid. Gray matter and white matter images were non-linearly registered to a custom template, derived from the training dataset (see below), using SPM12’s *DARTEL* toolbox (43). Using FSL’s *slicesdir* script^6^, default slices in sagittal, coronal, and axial directions were created for the segmented and normalized brain images. With these images, visual quality control was performed to ensure accurate image segmentation (Figure 1).

**Fig. 1.**
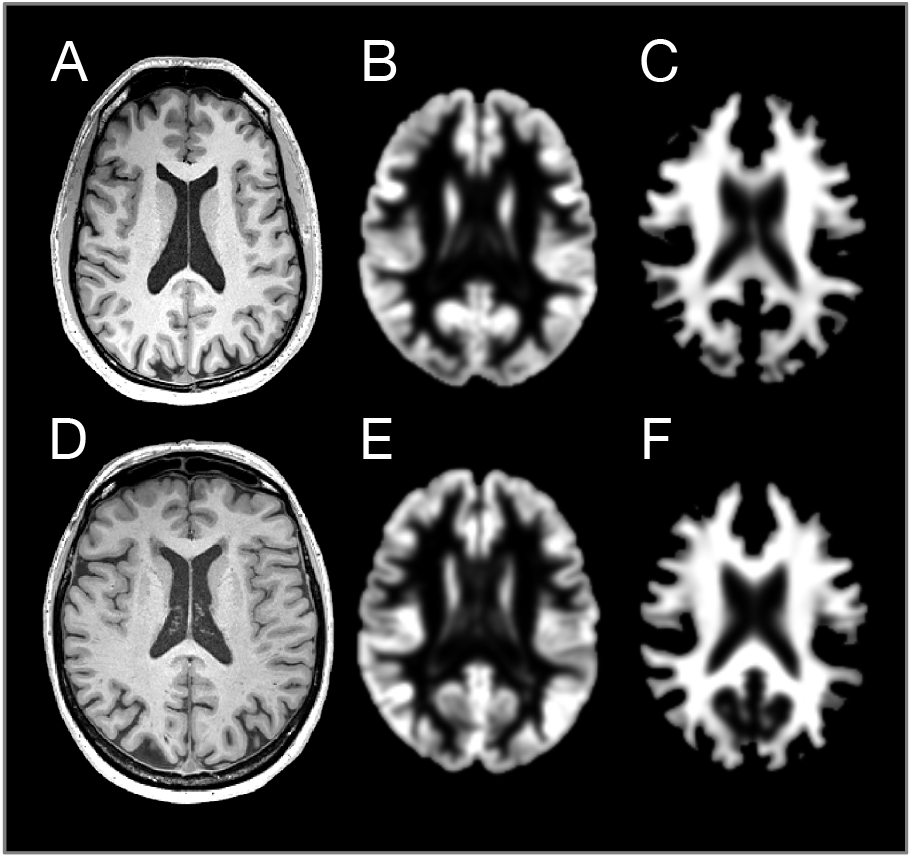
Original T1-weighted MR images (A, D), preprocessed gray matter (B, E), and preprocessed white matter images (C, F) of a 30-year-old (upper row) and a 66-year-old pain-free control participant (lower row). Segmentation was performed with SPM12’s *Segment* function. Original images are in individual space, preprocessed images were normalized to a brain template. Normalization was done with SPM12’s *DARTEL* toolbox. Preprocessed gray and white matter images were then vectorized and concatenated for brain age prediction.

The individual segmented and normalized images were loaded into R^7^(44) and vectorized. Gray and white matter vectors were combined. Brain age was then predicted with a Gaussian processes regression (45) using R’s *kernlab* (kernel-based machine learning lab) package^8^ (version 0.9-27) (46) and the previously learned regression model.

The *brainageR* model was trained on 2001 healthy individuals from several publicly-available datasets. The following studies contributed more than 200 T1-weighted images each: the Information Extracted from Medical Images (IXI) database^9^, the International Consortium for Brain Mapping (ICBM) study^10^ (47), and the Open Access Series of Imaging Studies (OASIS-1)^11^ (48). For details about all studies used for the training dataset, see Supplementary Information in Cole et al. 2017 (40) and 2018 (41). For further statistical analysis, brain age delta was calculated (predicted brain age minus chronological age).

### Statistical analysis

To check for normality, a Shapiro-Wilk test was performed. To check for equality of variances, Levene’s test was done. To compare the means of brain age delta in both groups, a traditional independent samples Welch’s t-test was performed (49); statistical significance was set at p < 0.05. We also performed a subgroup analysis with patients and controls ≥ 60 years of age using Welch’s t-test. In addition, a Bayesian independent samples t-test was calculated using the default Cauchy prior centered on zero and with r = 0.707 (50). For statistical testing, JASP^12^ version 0.92 was used (51). Individual brain age delta data^13^ and the results of statistical analysis^14^ have been published at the Open Science Framework.

## Results

### Demographics and clinical characteristics

Mean chronological age of patients with chronic pain was 53.0 ± 9.0 years (n = 59; 43 women). Mean age of the control group was 52.6 ± 9.0 years (n = 60; 44 women). In the group of chronic pain patients, mean pain duration was 15.9 ± 11.0 years (minimum: 1 year, maximum: 50 years).

In the ChroPain1 study, the primary pain diagnoses were degenerative spinal disease (n = 22), degenerative joint disease (n = 5), abdominal pain (n = 2), and fibromyalgia (n = 7). In the ChroPain2 study, the primary pain diagnoses were rheumatoid arthritis (n = 11), ankylosing spondylitis (n = 9), psoriasis arthritis (n = 2), and CREST syndrome (n = 1).

### Relationship between chronological age and brain age

Figure 2 illustrates the relationship between the chronological age of each participant and the brain age predicted by *brainageR* (blue: pain-free controls, red: chronic pain patients).

**Fig. 2.**
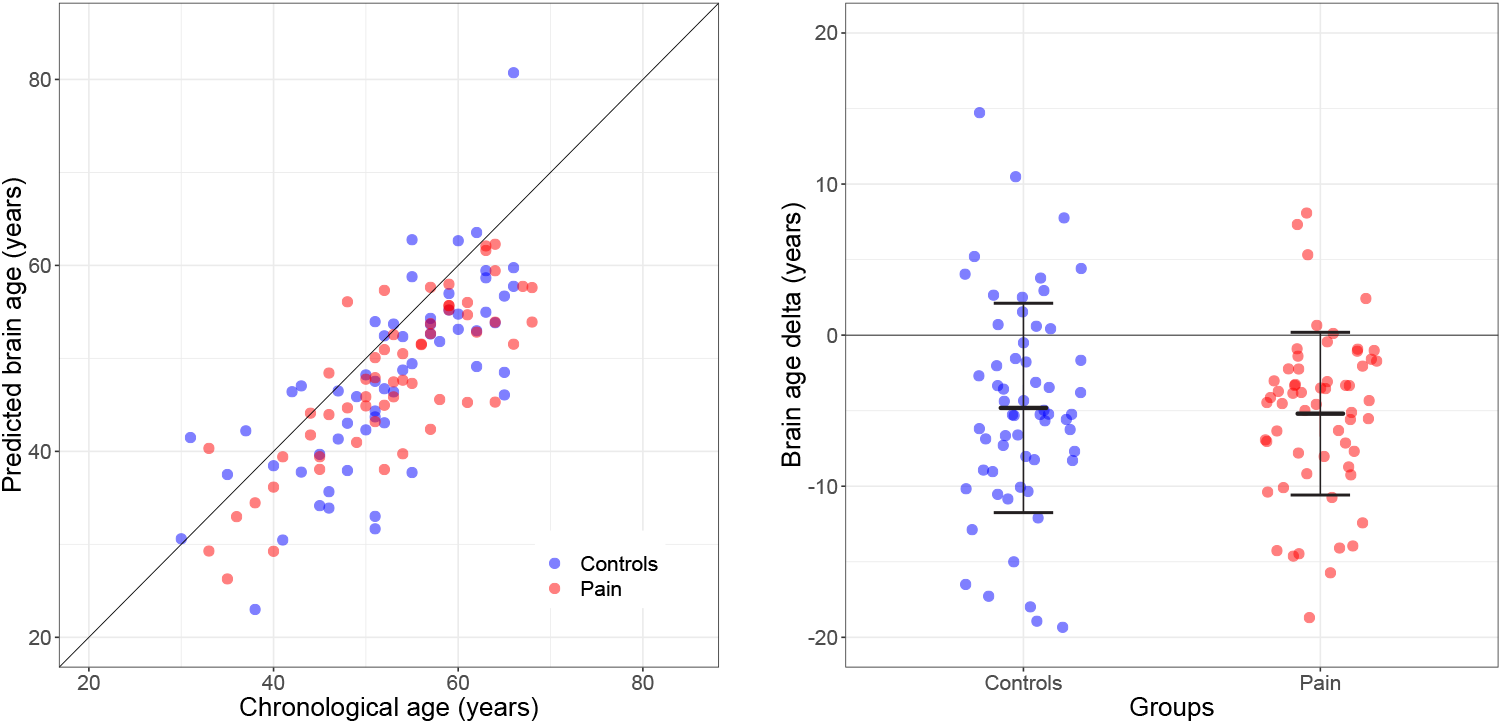
**A** | Relationship between chronological age and brain age predicted by *brainageR*. Blue circles represent pain-free control participants (n = 60), red circles represent patients with chronic non-cancer pain (n = 59). **B** | Brain age delta (predicted brain age minus chronological age) for pain-free controls (blue) and patients with chronic pain (red). The error bars symbolize mean ± standard deviation.

### Comparison of brain age delta between groups

Mean brain age delta ± standard deviation was −4.8 ± 6.9 years (minimum: −19.3, maximum: 14.7 years) for the control group and −5.2 ± 5.4 years (minimum: −18.7, maximum: 8.1 years) for the chronic pain group. The results of the Shapiro-Wilk test did not suggest deviation from normality in neither group (W = 0.983, p = 0.543 for the control group; W = 0.967, p = 0.104 for the chronic pain group). Levene’s test suggested equality of variances (F = 2.227, p = 0.138). Group means of brain age delta were not significantly different (Welch’s independent samples t-test; t(111.1) = 0.333, p = 0.74, Cohen’s d = 0.061). A Bayesian independent samples t-test indicated moderate evidence in favor of the null hypothesis (BF01 = 4.875; Figure 3).

**Fig. 3.**
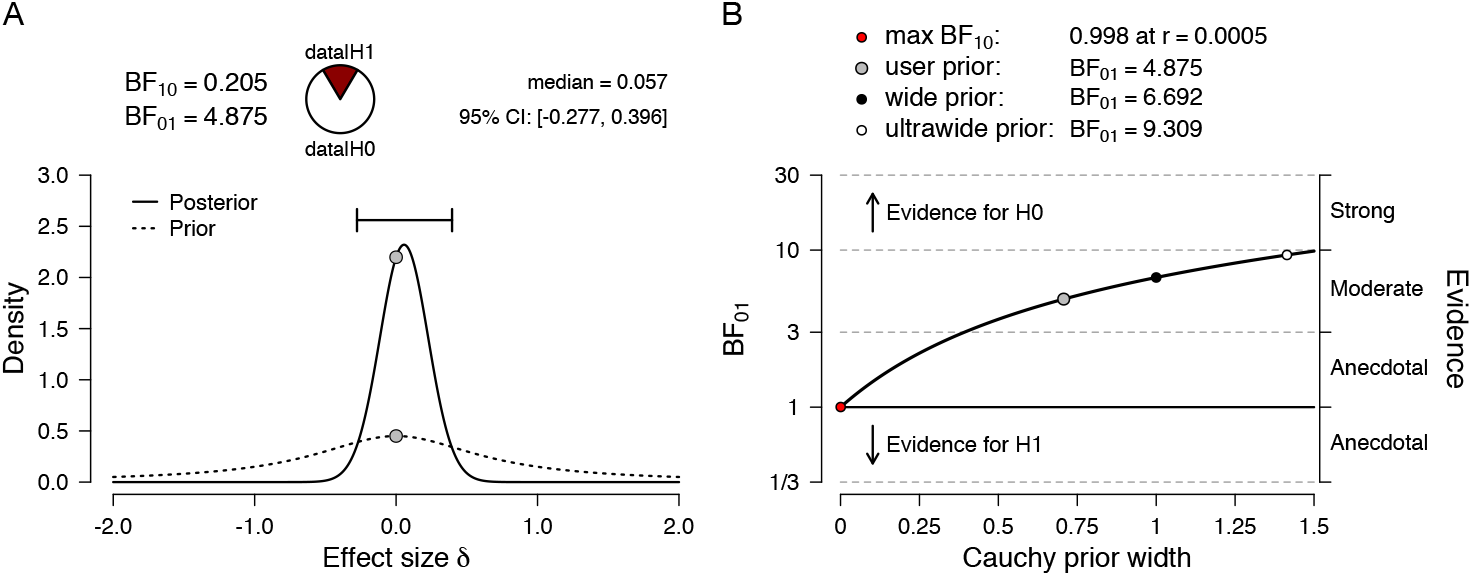
**A** | Bayesian independent samples t-test for the effect size δ. The dashed line illustrates the prior distribution (default Cauchy prior centered on zero, r = 0.707), the solid line shows the posterior distribution. The two gray dots indicate the prior and posterior density at the test value. The probability wheel on top visualizes the evidence that the data provide for the null hypothesis (H0: effect sizes are equal) and the alternative hypothesis (auburn, H1: effect sizes are different). The median and the 95% central credible interval of the posterior distribution are shown in the top right corner. **B** | The Bayes factor robustness plot. The plot indicates the Bayes factor BF01 (in favor of the null hypothesis) for the default prior (r = 0.707), a wide prior (r = 1), and an ultrawide prior (r = 1.414). All priors suggest moderate evidence for the null hypothesis, which is relatively stable across a wide range of prior distributions. Plots taken from JASP; the reporting of the results follows van Doorn et al. (52).

In a separate analysis with participants ≥ 60 years of age only, mean brain age delta was −6.2 ± 7.9 years in pain-free participants (n = 16) and −8.7 ± 5.7 years in chronic pain patients (n = 14; Welch’s t-test, p = 0.327, Cohen’s d = 0.362).

## Discussion

The present study of 59 chronic non-cancer pain patients and 60 healthy, age- and sex-matched controls with a mean age of 53 years provided no evidence for the hypothesis that chronic pain is associated with accelerated brain aging. In fact, a Bayesian independent samples t-test provided moderate evidence for the null hypothesis (i.e., group means are equal). Physiological aging is characterized by an inevitable – but also highly variable – decline in cognitive, motor, and sensory functions and also by changes of gray and white matter structure (53). In cross-sectional studies of healthy aging, voxel-based (54) and surface-based morphometry (55) have demonstrated a decrease of cortical gray matter density (56) and thickness (57), predominantly in prefrontal and temporal areas. Additionally, changes of white matter fiber tracts, as assessed by diffusion tensor imaging, have been investigated across the age span. In a longitudinal study on participants aged between 50-90 at baseline and with a follow-up after two years, a decline of white matter integrity was seen throughout the brain (58).

In recent years, several approaches have been developed to determine the degree of physiological or pathological aging of individual adults by predicting individual brain age (40, 41, 59–63). These studies have used individual structural MR images of the brain, large MRI training datasets of healthy individuals, and machine learning methods to compare individual brains with the training data (36). The results of these predictions have been shown to be accurate and reliable (36). Accelerated brain aging has been demonstrated in patients with Alzheimer’s disease (64), after traumatic brain injury (59), in HIV-positive individuals (65), and in psychiatric disease, such as schizophrenia, major depression, and borderline personality disorder (66). These methods are able to successfully predict conversion from mild cognitive impairment to Alzheimer’s disease (67). Moreover, in a group of 669 elderly participants with a mean age of 73 years (the Lothian Birth Cohort 1936 (68)) increased brain age delta was associated with decreased physical fitness (weaker grip strength, poorer lung function, slower walking speed), lower fluid intelligence, and increased mortality risk (41).

Of note, brain age delta results determined by these methods, including the software used in the present study, *brainageR*, are age-dependent; brain age of younger participants is over-estimated and brain age of older participants under-estimated (61). This inherent bias is driven by a regression toward the age mean of the training data set, as the error in the regression model is not orthogonal to age (61, 69). This explains why mean brain age delta was approximately −5 years in both groups of our study, where most participants were older than 40 years. Since we recruited a thoroughly age-matched control group for our chronic pain patients, this under-estimation of individual brain age does not weaken our conclusion that individual brain age is similar in chronic pain patients and in pain-free controls.

Very recently, Cruz-Almeida and co-workers published a comparison of brain age delta between 33 elderly individuals with chronic pain (mean age: 70.6 ± 5.5 years) and 14 individuals without chronic pain (mean age: 71.5 ± 7.3 years). While the mean predicted brain age was smaller than the chronological brain age in both groups (p = 0.592), participants with chronic pain had a larger brain age delta after adjustment for chronological age, sex, and exercise levels (ANCOVA, F [1,41] = 4.9; p = 0.033) (35). There are major differences between Cruz-Almeida et al.’s study and our study. Importantly, our study is characterized by a considerably larger sample size (119 vs. 47 participants), a younger age (the majority of participants is between 30 and 60 years of age), and a longer pain duration (15.9 ± 11 vs. 6.3 ± 8.8 years). Comparing only participants 60 years or older in our sample did not result in a significant difference of brain age delta between groups, neither, and thus did not provide evidence for the notion that chronic pain accelerates brain aging in seniors only.

Our results have important implications for the pathogenesis of structural alterations and, ultimately, cognitive deficits in patients with chronic pain. Our results suggest chronic pain does not induce wide-spread neural and glial degeneration, presumably the leading cause of age-related structural brain changes (70). Indirectly, our results support recent alternative models, mainly derived from animal research, of regional structural and functional brain alterations in chronic pain. These models imply network-specific (26, 71) neuronal loss triggered by excessive excitotoxic stimulation of NMDA receptors (72) and ongoning neuroinflammation associated with e.g. release of pro-inflammatory cytokines, such as interleukin 1beta (73, 74). In addition, synaptic plasticity, mainly in the primary somatosensory cortex (75) and the anterior cingulate cortex (76), as well as changes in the dendritic structure (77) are supposed to contribute to chronic pain-related structural remodeling. These models also suggest that the frequently observed cognitive deficits in chronic pain are the direct consequence of persistent nociceptive input, mediated by the aforementioned network-specific structural and functional changes, rather than the result of generalized accelerated aging of the brain.

## ACKNOWLEDGEMENTS

The authors wish to thank Dr. James Cole, King’s College London, London, UK, for his help with setting up the *brainageR* software and interpreting the results. The authors also thank Dr. Katharina Koch and Melanie Spindler M.Sc. for their active support of the ChroPain1 and ChroPain2 studies, including recruitment, data acquisition, and data analysis. We are grateful for the help of Dr. Markus Voglau and his team and of Mara Schier and the Selbsthilfegruppe Morbus Bechterew Oldenburg during recruitment for the ChroPain2 study. We also thank Katharina Grote MTRA and Gülsen Yanc MTRA for assisting with MRI data acquisition. We appreciate the helpful comments by Kathryn Mitchell on the final manuscript.

This work was supported by the Neuroimaging Unit, University of Oldenburg, funded by grants from the German Research Foundation (DFG; 3T MRI INST 184/152-1 FUGG and MEG INST 184/148-1 FUGG). The ChroPain1 study was also funded by an intramural grant from the School of Medicine and Health Sciences, University of Oldenburg, to C. B. (Forschungspool, 2015-1).

## Footnotes

1 https://www.rheuma-oldenburg.de

2 https://www.drks.de

3 https://github.com/rordenlab/dcm2niix

4 https://github.com/james-cole/brainageR

5 https://www.fil.ion.ucl.ac.uk/spm/software/spm12/

6 https://fsl.fmrib.ox.ac.uk/fsl/fslwiki/Miscvis

7 https://www.r-project.org

8 https://cran.r-project.org/web/packages/kernlab/index.html

9 https://brain-development.org/ixi-dataset

10 https://ida.loni.usc.edu

11 http://www.oasis-brains.org

12 https://jasp-stats.org

13 https://osf.io/2xd7h/

14 https://osf.io/dqpb5/

